# Characterizing dog communities adjacents to key shorebird sites in Chile

**DOI:** 10.1101/2025.01.20.630152

**Authors:** Gabriela Contreras, Ronny Peredo, Erik M. Sandvig, Betsabé Rodríguez, Benjamín Gallardo, Heraldo V. Norambuena, Fernando Medrano

## Abstract

Free-ranging domestic dogs can cause numerous negative impacts on public health, tourism, and the environment. Some of these impacts on wildlife are compounded and facilitated by human communities, for example through the provision of food, health care, and shelter, which altogether increase the dog’s life expectancy. Shorebirds are one of the groups adversely affected by dogs through nest predation, behavioral changes, and persecution during foraging and roosting periods. In Chile, dogs are a ubicutous threat in wetlands. In this study, we characterized the dog populations at the mouth of the Lluta River (Chacalluta area) and the Coihuín wetland (Coihuín-Chamiza area), both of international importance for shorebirds. We estimated the total abundance of dogs, their seasonal survival, and demographic and behavioral traits by photographic capture-recapture methods in eight surveys of 6.6 km^2^ for Chacalluta and 2.0 km^2^ for Coihuín-Chamiza. We estimated a population size of 167 individuals in Chacalluta and 505 individuals in Coihuín-Chamiza, a relatively low survival rate in both sites, a healthy body status in more than 80% of the dogs, small group sizes (1-2 individuals), and movements of 1,000 to 3,000 meters between locations. We report patterns associated with human ownership, indicating irresponsible pet ownership. This information will be the basis for developing public policies and evaluating strategies to lessen their impact on shorebirds, such as demographic control of the animal population, reduction of abandonment, and promotion of responsible pet ownership practices.

## Introduction

The domestic dog (*Canis lupus familiaris*) is the most common and widely distributed carnivore globally, with an estimated population of one billion individuals (Gompper 2014). This species has been crucial in the development of urban and rural communities for thousands of years through a close bond with humans as domesticated animals (Serpell 1991 and 2017; Gompper 2014). However, dogs cause a series of impacts in terms of public health, livestock, tourism, and the environment (Loss et al. 2013; Hampson et al. 2015; Lunney et al. 2011; Young et al. 2011; Doherty et al. 2017; Acosta-Jammet 2015, WCS 2019; Garde et al. 2013). Some of the critical factors in the impact of dogs on wildlife are directly related to human decisions, including the provision of food, veterinary care, and shelter, which in general increase dogs’ life expectancies (Morters et al. 2014; Svensson 2022). Human communities also allow dogs free transit, unsupervised wandering, and abandonment in wilderness areas (Gompper 2014; Villatoro et al.; 2016; Silva-Rodríguez et al. 2023). Internationally, dogs, cats, and rats have been identified as one of the main threats to biodiversity, causing the extinction of numerous species worldwide (Doherty et al. 2016 and 2017). The impacts that dogs can cause on wild species include direct predation, behavior changes, harassment, competition, and disease transmission (Ritchie et al. 2014; Banks & Bryant, 2007; Zapata-Ríos and Branch, 2016; Vanak et al. 2014; Weston and Stankowich, 2014; Furtado et al. 2016; Young et al. 2011; Acosta-Jamett et al. 2011). The World Organization for Animal Health (WOAH, 2023) defines two types of dogs— those with an owner, for which a person is responsible, and wandering dogs, which are any dog with or without an owner and without direct human supervision or control, including feral dogs—and it is generally described that the most significant impacts are caused by dogs that move without restriction (Gompper 2014).

Among the wildlife affected by the presence of domestic dogs, shorebirds are a group that it is particularly impacted by this threat. This group of birds belongs to the order Charadriiformes and is usually associated with open environments, especially wetlands. Most shorebirds spend at least part of their lives on tidal flats of estuaries or marine shoreline, and others use inland habitats such as grasslands, rivers, lakes, and lagoons (Martínez-Curci et al. 2021). The main impacts on this group of birds are predation and destruction of nests (eggs and chicks), behavioral changes, and disturbances during their feeding and resting periods (Burger 1986; Lafferty 2001; Lafferty et al. 2006; Weston et al. 2012). Dogs increase levels of disturbance and stress, and have been shown to be perceived as predators, even if they do not bother them directly (Lord et al. 2001; Glover et al. 2011; Drever et al. 2016; Murchison et al. 2016). While quantifying the effect of this threat is challenging, widespread and ongoing disruption to birds’ daily activities leads to significant energy costs, decreased fitness, and reduced reproductive success (Lord *et al*. 2001; Lafferty, 2001; Senner *et al,* 2017; Navedo et al. 2019; Gómez-Serrano, 2020).

The presence of dogs is one of the main disturbance agents recorded for shorebirds in coastal ecosystems in Latin America (Heredia-Morales 2019; Saiz-M et al. 2024), and Chile is no exception. Both resident and migratory shorebirds are also affected by the presence of dogs in their nesting, resting, and feeding areas (Cortés et al. 2019; Navedo et al. 2019; Díaz et al. 2024). Some species such as Killdeer (*Charadrius vociferus)*, Collared Plover (*Anarhynchus collaris*), Snowy Plover (*Anarhynchus nivosus*), American Oystercatcher (*Haematopus palliatus*), Black-necked Stilt (*Himantopus mexicanus*), Whimbrel (*Numenius phaeopus*), Hudsonian Godwit (*Limosa haemastica*) and Southern Lapwing (*Vanellus chilensis*) are threatened by dogs, causing them stress and limiting the development of critical activities for reproduction and migration (Aguirre 1997; Ortega-Solis et al. 2017; Oliveros et al. 2021).

In Chile, there are two key sites on the Pacific Flyway of the Americas where the presence of dogs has been identified as a potential threat to shorebirds: the Lluta River Mouth Wetland and the Coihuín-Chamiza Marine Wetland. Both sites are internationally recognized for hosting abundant populations of resident and migratory shorebirds (García-Walther et al. 2017; MMA-ROC-Manomet, 2023), and both have populations of shorebird species that have been classified under some category of threat under the national endangered species act, including the Snowy Plover (Vulnerable), Hudsonian Godwit (Vulnerable), and Collared Plover (Near-threatened). However, to date, there is little information quantifying the impact of dogs on the fitness on shorebirds at these areas. So far, the only information available indicates that dogs can predate on the nests of some resident shorebirds such as the American Oystercatcher and Snowy plover, both in Coihuín-Chamiza (Rodríguez 2023) and Chacalluta (Álvarez et al. in prep.).

This study aims to describe the dog population in the wetlands of the mouth of the Lluta River and the Coihuín marine wetland. Specifically, (i) we estimated population size and seasonal survival at both sites; (ii) we characterized the size of the groups; (iii) we characterized the body condition of individuals; and (iv) we characterized their movement in order to evaluate their potential effects on the shorebird populations inhabiting these wetlands. To do this, we carried out photographic censuses of free-ranging dogs between 2022 and 2023 in the localities adjacent to both wetlands.

## Materials and methods

### Study area

The study was carried out in localities adjacent to the wetlands of the mouth of the Lluta River, Region of Arica and Parinacota (18°24 ′ 32.21“S; 70°19 ′ 21.95”W) and the marine wetland of Coihuín-Chamiza, Region of Los Lagos (41°30’S; 72°50’W). Climatic conditions vary in both locations, with Arica (Chacalluta) having a desert climate and Puerto Montt (Coihuín-Chamiza) having a temperate climate. In the case of Arica (Chacalluta), the average temperatures fluctuate between 17.1°C for the winter months and 22.2°C for the summer months for the years 2022 and 2023. In Puerto Montt (Chamiza-Coihuín), winter temperatures average 6.4°C and summer temperatures average 14.8°C (Meteorological Directorate of Chile 2023).

The Lluta River is in the Chacalluta sector and has an area of 300 hectares, of which 30.64 are managed as a Municipal Nature Reserve and Nature Sanctuary. Meanwhile, the Coihuín marine wetland is adjacent to the towns of Coihuín and Chamiza, and covers an area of 1,765 hectares (Fig 1). For the purposes of this study, the monitoring area was defined at two zones: (a) priority area affected by the presence of dogs (area corresponding to 30.6 hectares in the Lluta River wetland and 1,765 hectares in the Coihuín marine wetland) plus the (b) area of influence associated with population settlements in the Chacalluta and Coihuín-Chamiza localities respectively (with a perimeter of 20.2 km and an area of 6.6 km^2^ in the Chacalluta area, and a perimeter of 15.3 km and an area of 3.1 km^2^ in Coihuín and Chamiza, Fig 1). Henceforth, the study areas will correspond to Chacalluta (Arica) and Coihuín-Chamiza (Puerto Montt).

**Figure 1.**
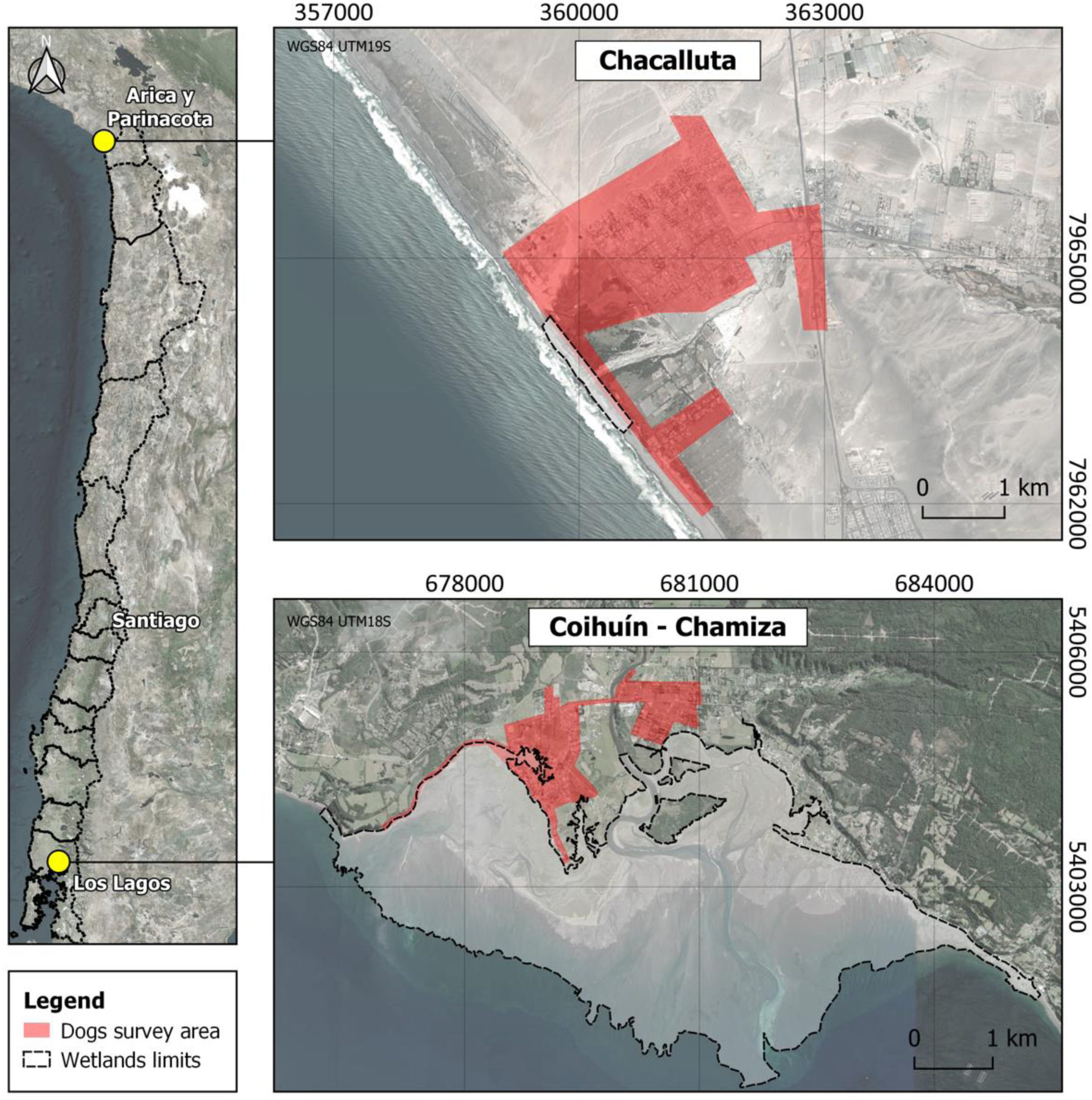
Study areas in the wetland at the mouth of the Lluta River (Chacalluta) and the marine wetland of Coihuín (Coihuín-Chamiza).

Both wetlands are classified as Important Bird Areas (IBAs), are important sites for the Western Hemisphere Shorebird Reserves Network (WHSRN) and are recognized in the Pacific Flyway of the Americas Shorebird Conservation Strategy. The mouth of the Lluta River is managed at the municipal level and presents an important set of threats associated with its proximity to the city, tourism, and recreational activities. The marine wetland of Coihuín includes an extensive tidal plain, with surrounding sectors that are increasingly urbanized, and with threats related to human population growth (Delgado & Cursach 2021). Residents of this site include members of two indigenous Mapuche Lafkenche communities, who are seeking the designation at the national level of the area as an Indigenous Peoples Marine Coastal Space (ECMPO).

### Surveys

The dog population was estimated by photographic capture-recapture, surveying 667 hectares for Chacalluta and 308 for Coihuín-Chamiza, adjusted to the city’s road network, a method successfully used in Ushuaia by Arona & Shiavini (2023). Eight vehicle surveys were carried out during daylight hours, on days without rain, with an approximate duration of 4 hours, covering the entire sampling area. We photographed each dog that transited without restriction and recording its body condition (body condition, skin health or presence of lesions) as well as the presence of accessories (collar, clothing). Each photograph was georeferenced based on the synchronization of the capture time of the cameras and the GPS. The routes were carried out at a speed of approximately 15 km/h, allowing enough time to classify the state of well-being and characteristics of the dogs recorded.

Data recording was carried out on board a vehicle with wide visibility, while one or two volunteers recorded the dogs observed on each side of the vehicle through photographs, increasing the probability and efficiency of detection (Arona & Shiavini 2023). This methodology was chosen because it does not require physically capturing dogs to mark them and assumes that (1) the capture of the animal does not affect its subsequent probability of recapture, (2) on a sample occasion all animals have the same probability of being registered, (3) there are no additions (through births or immigration) or losses (mortality or emigration), and (4) identifying marks are not lost during the study (Arona & Shiavini 2023).

Dogs that were freely roaming unrestricted on public roads were considered, as well as dogs that were inside dwellings but had free passage to leave (unrestrained dogs and dogs in dwelling without an effective perimeter fence for exit). Likewise, in order to have information on the characteristics of roaming dogs, each dog was registered on a spreadsheet and classified according to a modification of Hiby & Hiby (2017) categorization: (a) visible well-being, through body condition index on a scale of 1 to 3 points, based on body fat coverage (underweight, ideal weight, and overweight, respectively) in addition to the visible condition of the skin (coat health or presence of lesions) or difficulties in movement, and (b) collar ownership, i.e., possession of collars or indicator accessories of dogs with an owner (Arona & Shiavini 2023). The route was recorded with GPS using a smartphone application that allows recording geographical coordinates and other data (Locus Map).

The first census represented the “capture” of the dogs observed, while the records of subsequent monitoring represented the “recaptures” of previously captured dogs as well as “new” dogs. The photographs were analyzed individually for each route and were compared between different samples. Phenotypic characteristics were used to identify the animals individually, verifying coincidences of occurrence between the censuses. Dogs that had already been seen in previous sampling (recaptured) and those that had not been previously found were identified. Each dog was categorized with an identification code for further analysis.

To determine dog abundance at Chacalluta and Coihuín-Chamiza, eight surveys were conducted at each site during June, July, November, and December 2022, June and July 2023, and January and February 2024. These eight replicates were conducted during two main periods (austral winter and summer), photographing and individually identifying dogs sighted during a predetermined transect (Arona & Shiavini 2023).

### Data analysis

Identifying dogs at the individual level allowed us to create a database with capture-recapture histories for everyone, to which we applied a ’robust design’ model (Firth & Bennett 1998) using the *rCapture* package (Baillargeon & Rivest 2007; Rivest & Baillargeon 2022) in R (R Core Team 2024). This model assumes that the population remains closed (no births, mortality, immigration, or emigration) within a survey but that it is open between surveys. This allows estimating the abundance and recapture rate within each major period and the survival rate between the major periods. An equal recapture rate was assumed for all individuals detected within a period.

To analyze the locations where each of the dogs was found during the censuses and observe the phenomenon in a more comprehensive way, we used a geospatial tool present in the QGIS software called “Heat Map (Core Density Estimation)”, which is used in spatial analysis to visualize the distribution of specific events on a map (QGIS 2024).

To complement the results of this study and identify dog movements, we used information collected through tracking cameras in the Coihuín wetland at 11 shorebird nests during the breeding seasons of 2021 and 2022 (Rodríguez 2023). From these data, we analyzed the distance traveled by dogs from shorebird breeding areas to surveyed sites.

Finally, to determine the dog:humans ratio, we used the national surveys by the National Institute of Statistics (INE) for 2017. In the case of Chacalluta, we used data from Villa Frontera and La Ponderosa. In the case of Coihuín, we used the polygons adjacent to the wetland and those considered within the study.

## Results

A total of 175 photographs were taken with 114 dogs identified for Chacalluta and 265 photographs with 267 dogs identified for Coihuín and Chamiza.

### Dog population size and survival rate

A total abundance of dogs of 505.14±33.38 individuals was estimated for Coihuín-Chamiza and 167.54±14.00 individuals for Chacalluta (Fig 2A) for the study period. In the case of Coihuín-Chamiza, a higher abundance was estimated for the summer of 2023 (491.70±58.69), and a lower abundance during both winters and the summer of 2024 (295±64.81, 275±31.31, and 267±57.90). In the case of Chacalluta, no pattern associated with seasonality was observed, although an apparent decrease was observed over the course of the surveys (120.00±19.99, 126.19±16.78, 100.88±11,47, and 78.00±36.06). Recapture rates were lower during summer than winter at Coihuín-Chamiza (Fig 2B; 0.25±0.08-0.36±0.16 and 0.58±0.09-0.65±0.07, respectively). While at Chacalluta there was a lower recapture rate in summer than winter between the first two surveys, but an overlap between seasons in the third and fourth survey (Fig 2B; 0.22±0.03-0.38±0.08 and 0.39±0.08-0.42±0.05, respectively). In Coihuín-Chamiza similar survival rates were estimated for winter 2022 to summer 2022/23 (0.99±0.14) and winter 2023 to summer 2023/24 (0.93±0.22), and a much lower survival from summer 2022/23 to winter 2023 (0.56±0.08), while in Chacalluta there was an apparent decrease in the survival rate between survey periods (Fig 2C; 0.82±0.13, 0.80±0.24, 0.58±0.29).

**Figure 2:**
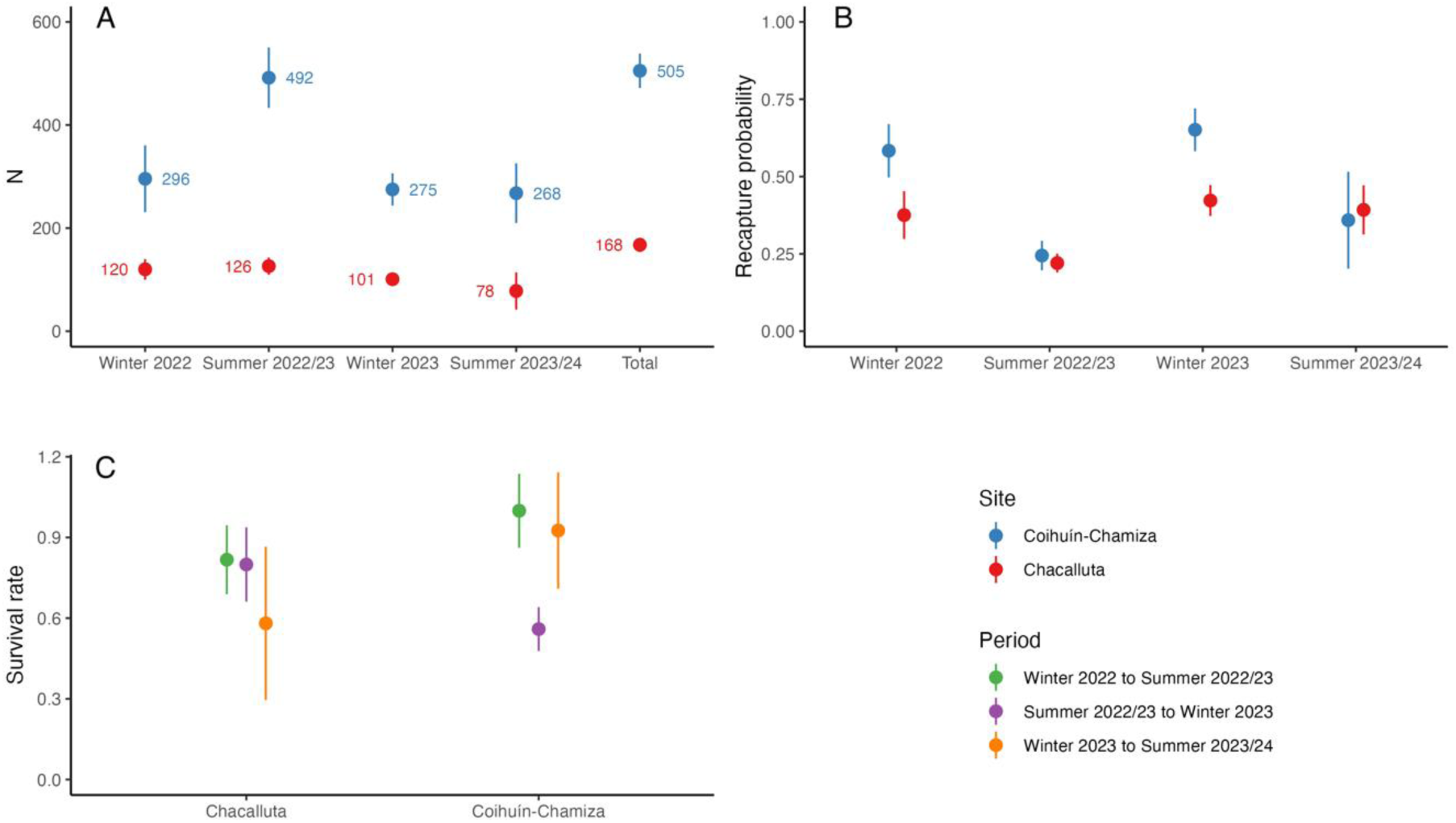
A) Dog abundance estimates for each survey period and overall for both study sites; B) Dog recapture rates for each survey period; and C) Survival rate estimates between each survey period derived from a ‘robust design’ capture-recapture model.

### Dog:inhabitants ratio

In Coihuín, the population was reported at 1,686 people in 197 hectares, resulting in a population density of 8.56 inhabitants/hectare and a dog:human ratio of 1:3.3, or 1 dog for every 3.3 inhabitants. For Chacalluta, with data from the populations of Villa Frontera and La Ponderosa, the population was 940 people, with a density of 3.34 inhabitants/hectare, and a dog:human ratio of 1:5.6.

### Characterization of the dog population

Regarding the origin of the detected dogs in the Coihuín-Chamiza site, almost 60% of the dogs recorded were on public areas and the rest were detected within private properties. In the case of Chacalluta, almost all dogs (97.9%) were observed in public areas (Fig 3A). The frequency of dogs with injuries was less than 5% in both sities (Fig 3B).

**Figure 3.**
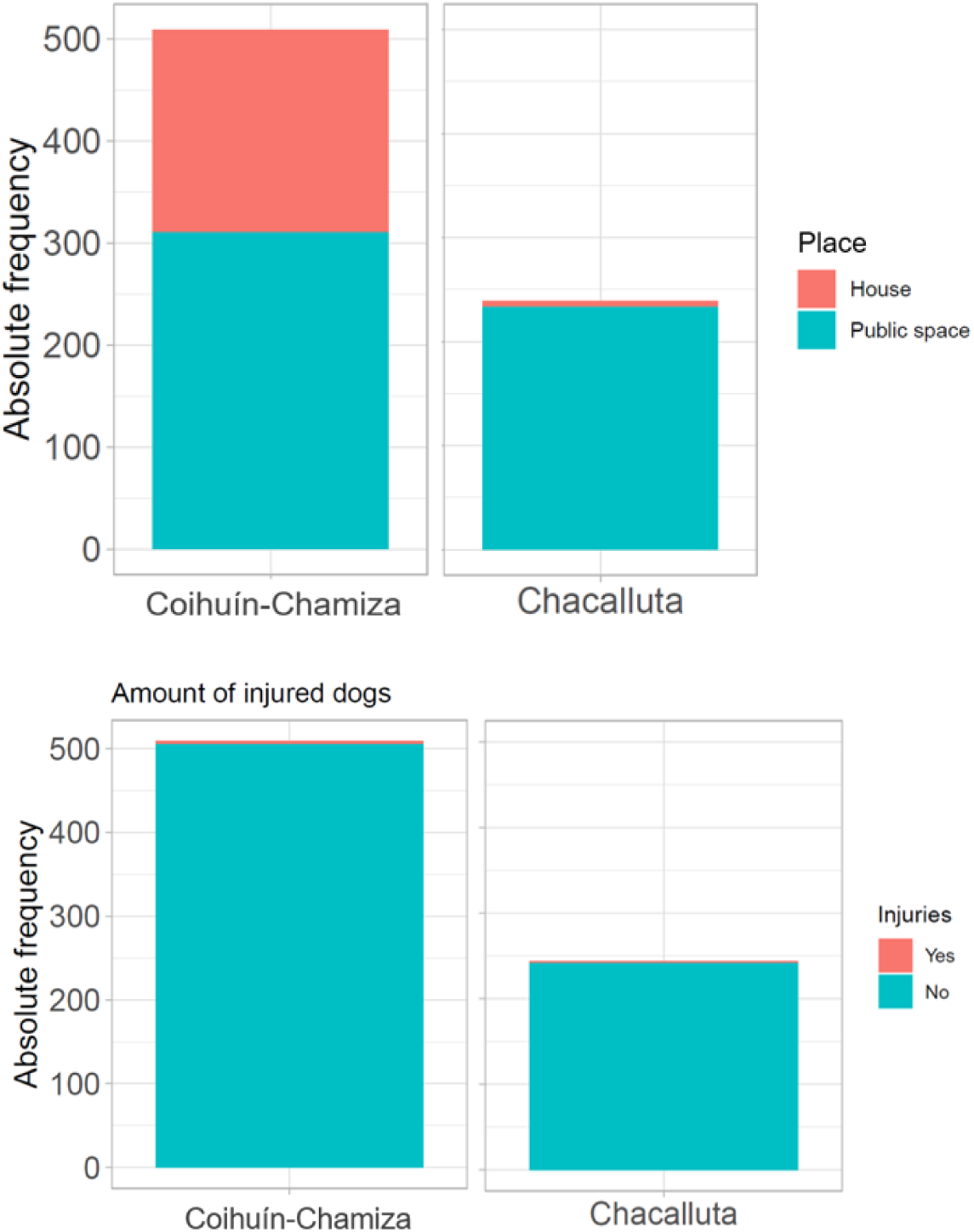
A) Absolute frequency of origin of dogs for both sites, B) Absolute frequency of injuries in dogs for both sites.

Most dogs presented an ideal body condition at both sites (97,3%), and very few individuals were underweighted (2,3%) or overweighted (0,3%, Fig 4A). Most detections were of solitary individuals (52,7%), indicating that most dogs move on their own in these areas. We also observed 43.7% small groups (<5 individuals) in Coihuín-Chamiza and 47.5% in Chachalluta. In rare instances we observed larger groups, with up to 9 dogs in Chachalluta and 19 individuals in Coihuín-Chamiza, suggesting a greater congregation of dogs in specific areas, but as isolated events with respect to the observed pattern (Fig 4B).

**Figure 4.**
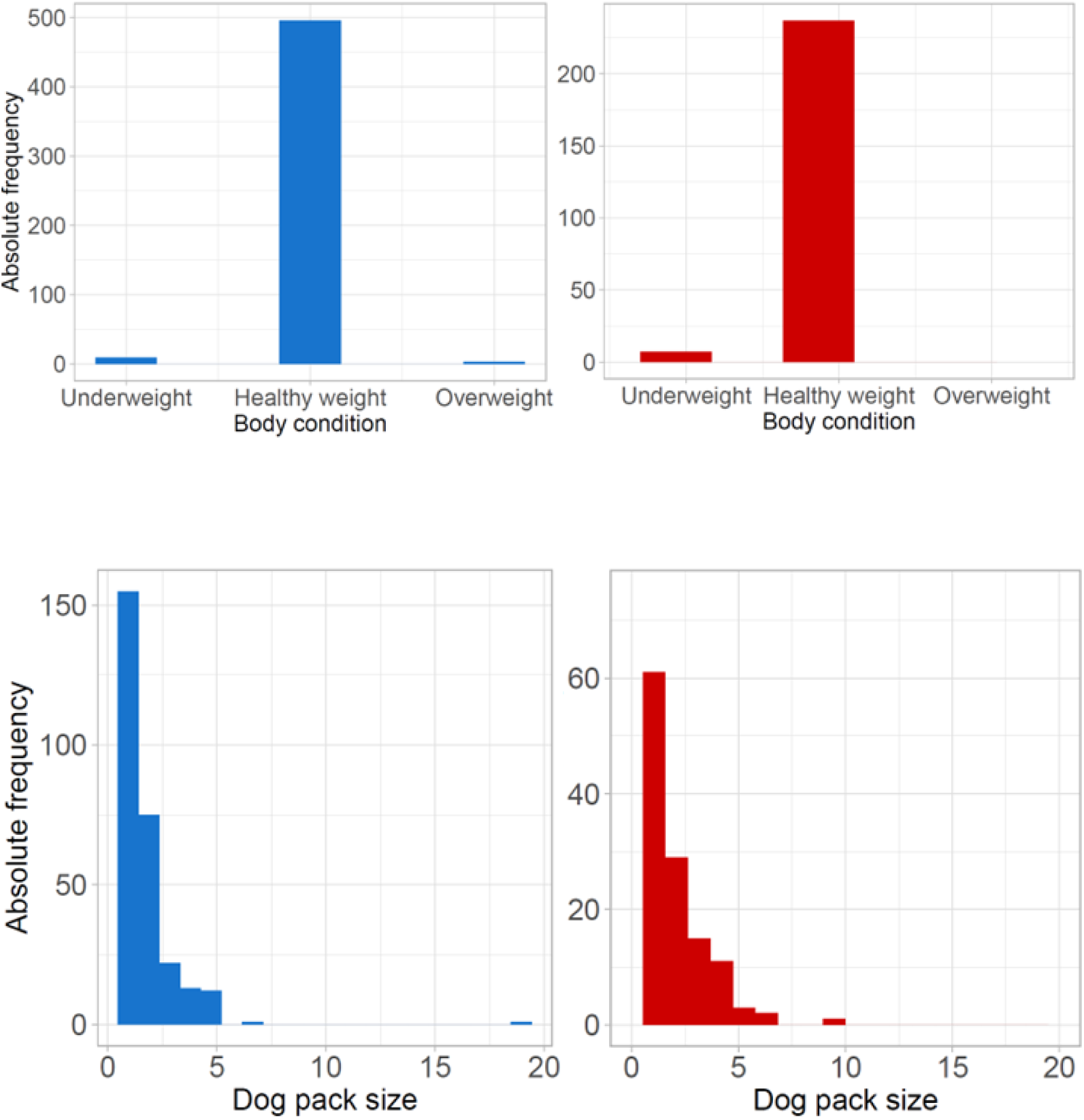
A. Frequency of body condition in unrestricted dogs at both sites. B. Size of packs of dogs without restriction of movement at both sites. (Blue bars represent Coihuín-Chamiza values, while red bars represent Chacalluta values).

### Range of movement

Most dogs (>X%) moved less than 1,000 m at both sites, and in Chacalluta, X% moved less than 200 m. Movements between 1 and 2 km were observed only in X% of the individuals, and movements >2 km occurred only with X% of the dogs, suggesting that large-distance movements are rare in the studied areas (Fig 5A). We only detected 7 dogs with the camera-traps located at shorebird nests within the wetlands, all of them with movements of 800-850 meters from their detection sites in the urban areas (Fig 5B).

**Figure 5.**
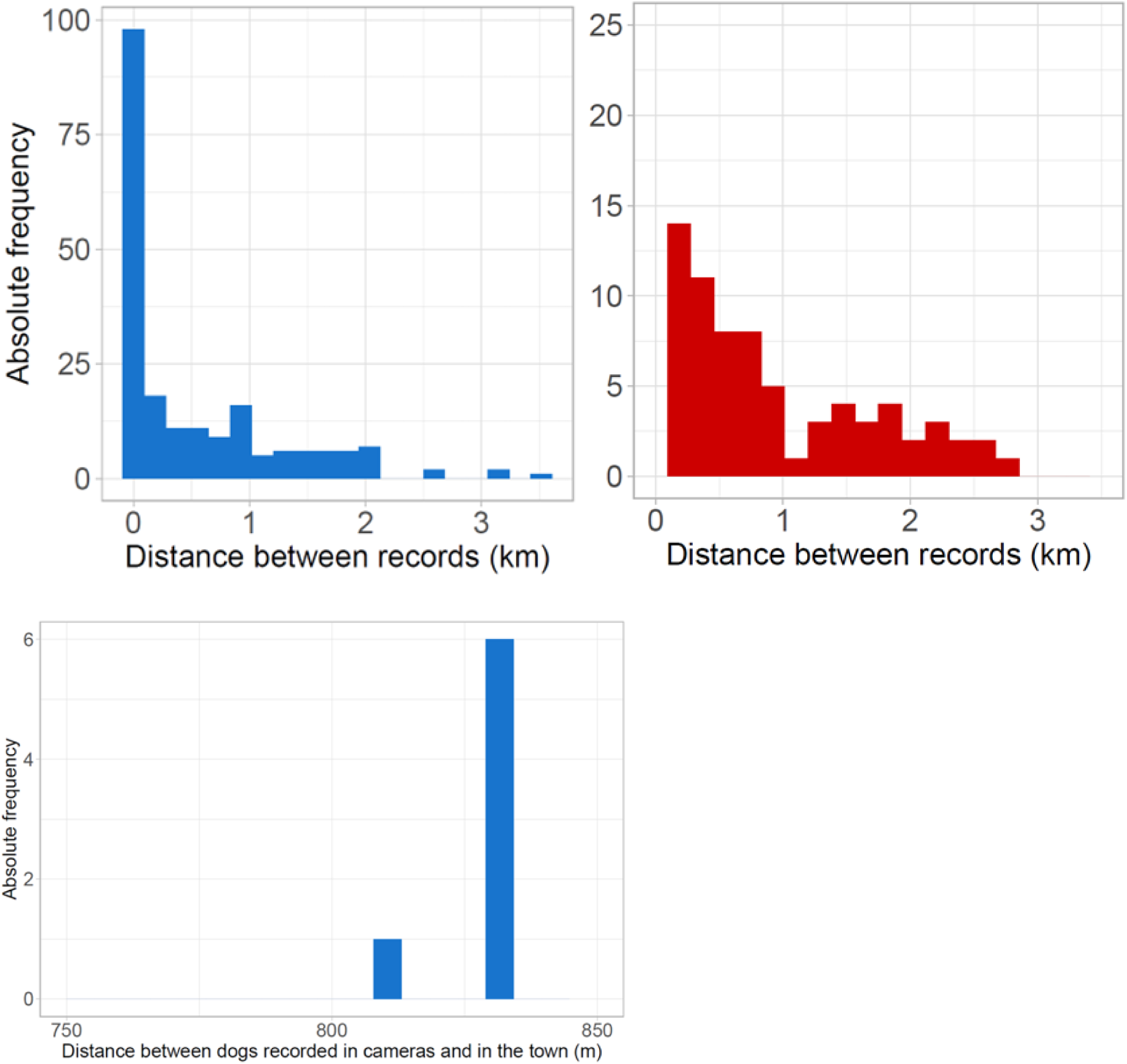
A. Displacements of census dogs in Chacalluta and Coihuín-Chamiza. B Distance between camera-trap registration of dogs and the urbanised area of Coihuín-Chamiza. (Blue bars represent Coihuín-Chamiza values, while red bars represent Chacalluta values).

### Spatial patterns

With the results of heat maps, in Chacalluta we observed a greater clustering of dogs in the sector of Villa La Frontera, an area with the highest population influx (Fig 6A). When analyzing the surveys by season, we observed that the records during the winter months move a little further away (between X and Y m) from the urban areas, unlike what happens in summer, where they are concentrated in more specific points near town (Fig 7A, 7B).

**Figure 6.**
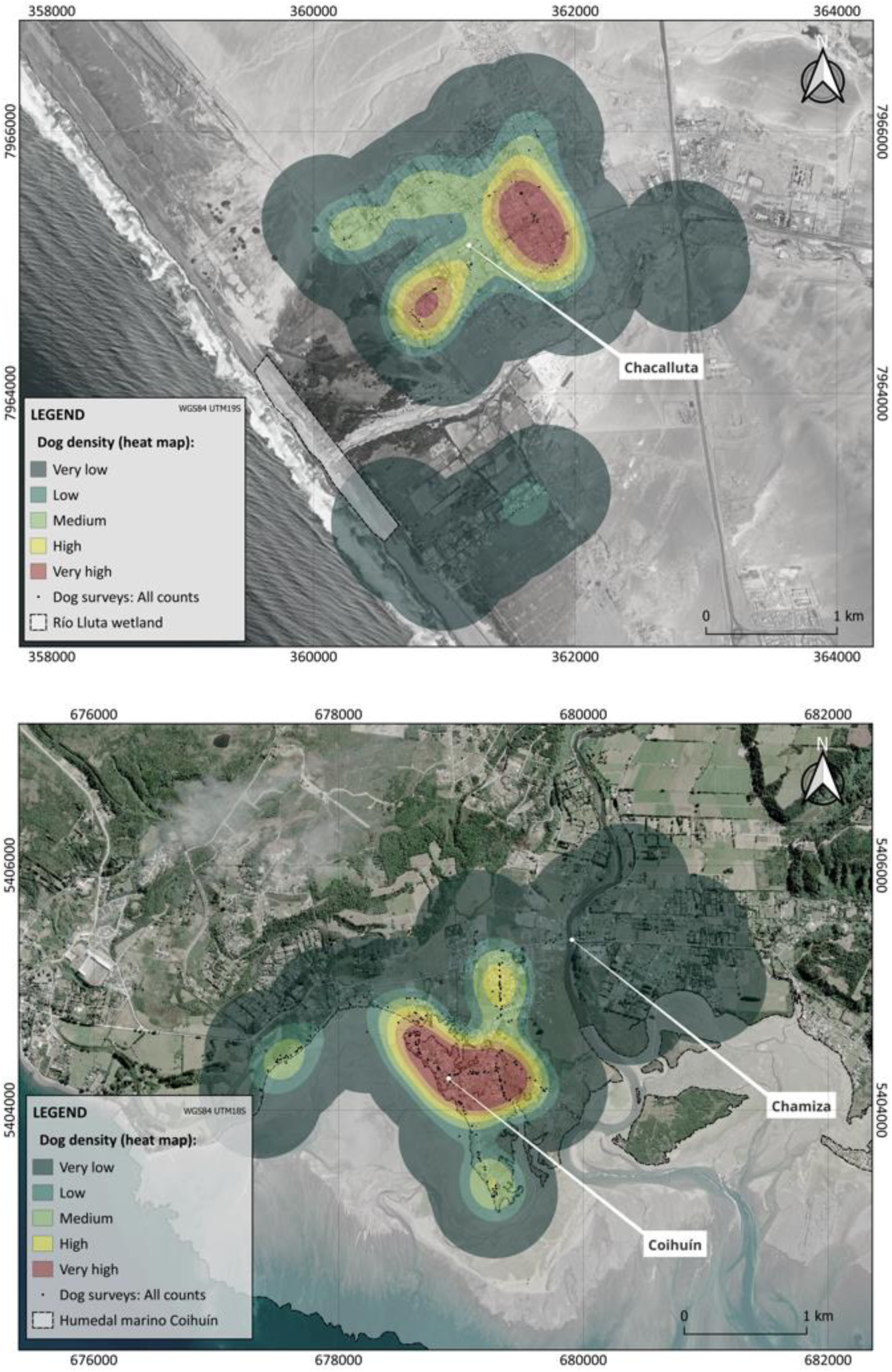
A. Map of heat areas in Chacalluta area with all data. B Map of heat areas in Coihuín-Chamiza area with all data

**Figure 7.**
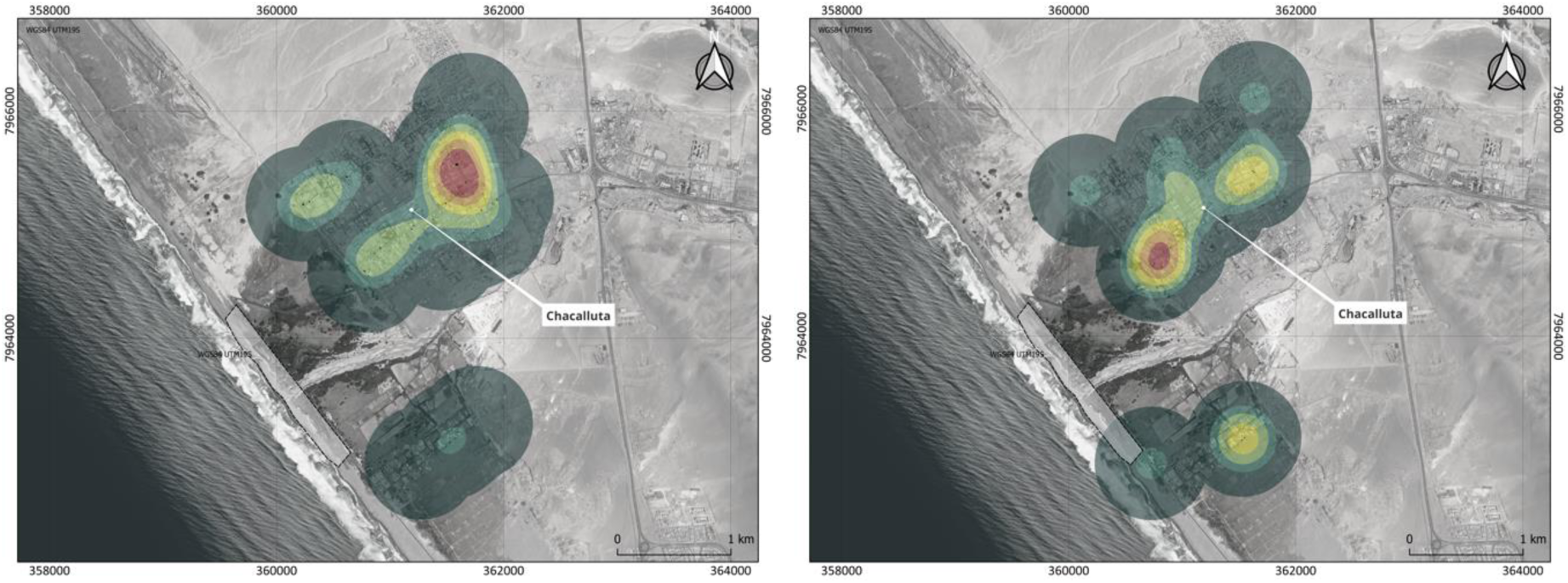
A Map of heat areas in the Chacalluta area during winter 2022. B Map of heat areas in the Chacalluta zone during summer 2024.

A similar pattern was observed in Coihuín and Chamiza, where the highest concentration of dogs occurs in places with higher population density, associated with the urban locality of lower Coihuín (Fig 6A). During the winter months, the dog records were concentrated on the urban area, while in the summer months, dogs were dispersed farther away (>X m) from the concentration of dwellings (Fig 8A, 8B).

**Figure 8.**
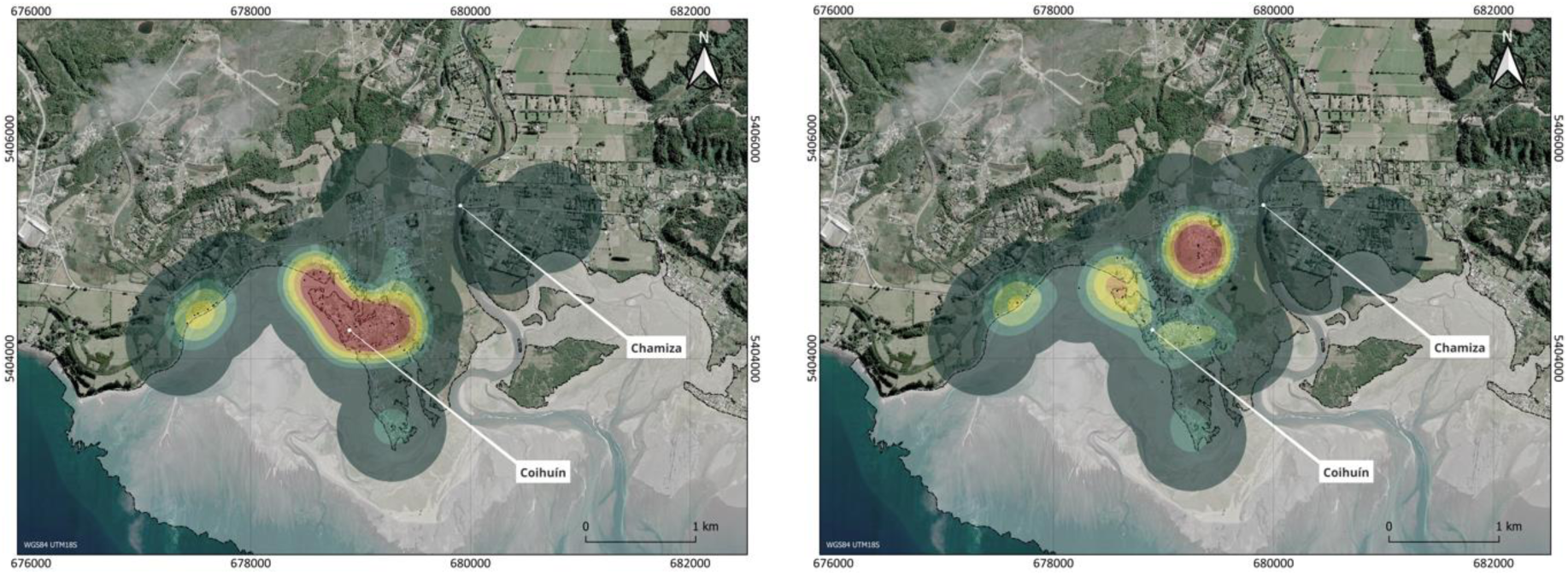
A Map of heat areas in the Coihuín-Chamiza area during winter 2023. B Map of heat areas in the Coihuín-Chamiza zone during summer 2022.

## Discussion

In this study we found a high abundance of dogs in both sites, with larger numbers within or in the proximity of urban areas. In the case of Chacalluta, no seasonal patterns were observed, which may be because temperature fluctuations do not vary significantly between seasons (from 15°C to 22C, Meteorological Directorate of Chile 2023). In the case of Coihuín, there were seasonal variations (5°C a 15°C, Meteorological Directorate of Chile 2023)., with higher abundances during summer than in winter, suggesting that dogs shelter during adverse weather conditions which may reduce detectability for ‘recapture’. Also, we found low dog:human ratios in Chacalluta and in Coihuín-Chamiza, for every 3 and 6 humans there is 1 dog respectively. These results possibly underestimate the true ratio, due to the low detectability of dogs found indoors or in non-visible parts of roads or yards. Considering these results, both cases are lower than those published by the Undersecretariat of Regional and Administrative Development of the Ministry of the Interior and Public Safety (SUBDERE), through the Responsible Pet Ownership Program (PTRAC), which establishes a ratio of 2.5 dog:inhabitant in urban areas and 1.7 in rural areas (SUBDERE, 2022).

Both sites present conditions that drive different pressures on the movement and survival of dogs. The fact that in Chacalluta there is a similar population size and recapture rate in winter as in summer can be interpreted as dogs being able to roam freely throughout the year due to stable weather conditions, impacting wetland shorebirds both in the breeding and non-breeding season. On the other hand, in Coihuín-Chamiza the dogs seem to be more sheltered in winter, but during the austral summer there would still be pressure on shorebirds, a period that coincides with the breeding season and when significant numbers of trans-equatorial migrants visit the place (e.g. Hudsonian Godwit).

At both sites, we observed a low survival rate for a relatively long-lived animal, with global estimates ranging from 10 to 12 years (O’Neill et al. 2013). However, adverse survival conditions coupled with irresponsible pet ownership could be a factor (food availability, disease, accidents; Astorga et al. 2022). These results could be underestimated due to relatively low detectability (recapture rate).

In addition, at both sites, the dogs were in groups of less than five individuals, indicating that in the area, they do not need to form large groups to survive and move. This could indicate that most dogs are under irresponsible ownership and are not feral dogs (wanderers that do not depend on humans and reproduce), which maintain certain survival behaviors. Typically, wandering dogs (those that roam freely, dogs that have been lost or abandoned) form large groups when resources are scarce and dispersed, and under the presence of predators, to increase their chances of survival (Boitani et al. 2017). Our results suggest that these conditions are not present in our study areas.

It could be suggested that dog detections at both sites are likely concentrated in areas of lower economic income, particularly around irregular settlements. Conversely, in Coihuín-Chamiza, areas with higher economic income (subdivisions) appear to have a lower abundance of wandering dogs. Although assessing the relationship between dog presence and socioeconomic status was not the study’s objective, the results would indicate that higher abundance occurs in poorer areas, which would be valuable to evaluate in future research. Additionally, it would be pertinent to consider the perception that residents have regarding the dog issue in relation to public health. This observation may align with global patterns, indicating that regions with greater economic resources tend to experience lower human densities, which could, in turn, lead to reduced dog densities. It is also assumed that in areas with limited economic resources, maintaining protective perimeters (e.g., fences, barriers) may be more challenging (Astorga et al. 2022; Silva-Rodríguez et al. 2023). Likewise, more frequent compassionate behaviors are described in lower-income neighborhoods when feeding stray dogs (Silva-Rodríguez et al. 2023).

Dogs from both sites were mainly detected on public roads or in houses without access restrictions, which coincides with what has been found in the literature, where dogs that live in rural areas usually have an owner or holder (Silva-Rodríguez et al. 2023), and free-roaming dogs are not identified as feral (WOAH, 2023). This could influence the condition of the dogs, as we found that most dogs were in good body condition and with a very low percentage of visible lesions. This situation could indicate that most dogs have some degree of human dependence (supervision or maintenance), possibly including food or other features such as shelter (Astorga et al. 2022; Silva-Rodríguez et al. 2023).

Our results on low distance movements may indicate that dogs in these areas do not need long journeys to obtain sufficient food or shelter, and that those responsible for the pets are located near the sighting locations. These observations are consistent with other studies where dogs in properties, but with access to the outdoors, concentrated their activities at a short distance from the owner’s home (Sepúlveda et al. 2015; Silva-Rodríguez et al. 2023) and when incursions farther from urban areas occur, the movements mainly follow human modified landscapes (Sepulveda et al. 2015).

A study published by Sepúlveda et al. (2015) analyzed the movements of dogs in rural areas of southern Chile using GPS radio collars. The results showed that dogs were primarily located within 200 meters of homes but exhibited movement patterns with distances ranging from 0.5 to 1.9 kilometers. This information is consistent with the data obtained in this study, where dogs recorded during the census were compared with those captured by camera traps (Rodríguez 2023) at American Oystercatcher breeding sites. It was observed that dogs moved less than 850 meters, suggesting that they tend to remain in urban areas. This implies that the dogs impacting the reproductive success of American Oystercatchers likely originate from areas close to these breeding sites. Furthermore, the evidence from Rodríguez highlighted that the presence of dogs is one of the main synanthropic impacts on the reproductive success of the American Oystercatcher at the breeding site in the Coihuín wetland, emphasizing the need for targeted conservation efforts in these critical areas.

To develop responsible pet ownership programs, the first step is to assess human perception of the conflict, demographics of companion animals, and local needs and resources within a specific area (Garde et al. 2022). Local involvement and community support are critical to the success of management programs, as integrating human health and animal welfare objectives into canine management can achieve greater community buy-in rather than focusing solely on species conservation (Doherty 2017). Crespin and Contreras-Abarca (2024) described that socioeconomic inequality precedes canine density, where poverty levels can influence the density of dogs that move without restriction. Therefore, it is critical to consider the priorities and needs of the inhabitants when planning actions to address this issue and evaluate and prioritize the diversity of environmental and socioeconomic factors at the local level.

Main measures for controlling stray dog populations include improving movement restrictions and promoting adequate confinement (OIE 2009, Astorga et al. 2022). Programs aimed at controlling loose dog populations should enhance owners’ awareness of their dogs’ impact, as these dogs may pose a threat to the public and wildlife, be disruptive, or disturb the community. Responsible pet ownership programs and the development of social norms and sanctioning policies can effectively promote confinement (Astorga et al. 2022). In cases where dogs are generally in good condition, reflecting a population’s interest in animal welfare, it is recommended to collect information about people’s perception of the stray dog conflict. This allows for outlining strategies, such as reducing attacks if dogs are seen as a risk or promoting fencing improvement campaigns if concerns are associated with pet welfare.

The study underscores the concerning prevalence of free-roaming dogs with owners in Chacalluta and Coihuín-Chamiza, highlighting potential threats to wildlife, particularly shorebird populations. While the findings emphasize the need for targeted interventions to promote responsible pet ownership, it is important to recognize the diverse realities that may exist across different local contexts. These actions should be informed by thorough assessments of local needs and resources, ensuring that interventions are contextually relevant and effectively address the unique challenges faced by each community. Additionally, further studies are necessary to comprehensively evaluate the impact of dog presence on shorebird behavior, distribution, and reproductive success. This would provide a more robust foundation for developing targeted conservation and management strategies to mitigate the negative effects on shorebird populations.

## Acknowledgments

To Sharon Montecino and Ivo Tejeda for their help in applying for and obtaining funds from the USFS project “Protection of shorebirds through disturbance management at two key sites in Chile.” To Emiliano Arona for presenting his work conducted in Ushuaia, Argentina, which provided valuable insights and facilitated the development of the USFS project. To Osvel Hinojosa for supervision and editing of the manuscript.

